# SARS-CoV-2-specific memory B cells can persist in the elderly despite loss of neutralising antibodies

**DOI:** 10.1101/2021.05.30.446322

**Authors:** Anna Jeffery-Smith, Alice R Burton, Sabela Lens, Chloe Rees-Spear, Monika Patel, Robin Gopal, Luke Muir, Felicity Aiano, Katie J Doores, J. Yimmy Chow, Shamez N Ladhani, Maria Zambon, Laura E McCoy, Mala K Maini

## Abstract

Memory B cells (MBC) can provide a recall response able to supplement waning antibodies with an affinity-matured response better able to neutralise variant viruses. We studied a cohort of vulnerable elderly care home residents and younger staff, a high proportion of whom had lost neutralising antibodies (nAb), to investigate their reserve immunity from SARS-CoV-2-specific MBC. Class-switched spike and RBD-tetramer-binding MBC with a classical phenotype persisted five months post-mild/asymptomatic SARS-CoV-2 infection, irrespective of age. Spike/RBD-specific MBC remained detectable in the majority who had lost nAb, although at lower frequencies and with a reduced IgG/IgA isotype ratio. Functional spike/S1/RBD-specific recall was also detectable by ELISpot in some who had lost nAb, but was significantly impaired in the elderly, particularly to RBD. Our findings demonstrate persistence of SARS-CoV-2-specific MBC beyond loss of nAb, but highlight the need for careful monitoring of functional defects in RBD-specific B cell immunity in the elderly.

**One sentence summary:** Circulating class-switched spike and RBD-specific memory B cells can outlast detectable neutralising antibodies but are functionally constrained in the elderly.

## Introduction

The human coronavirus SARS-CoV-2 has had a particularly devastating impact on the elderly, who are at much greater risk of morbidity and mortality.^1,2^ Understanding the nature of a successful immune response in those who have avoided these outcomes and cleared SARS-CoV-2 after a mild infection, despite advanced age, is key to protecting this vulnerable group in the future. Whether older survivors of SARS-CoV-2 infection are able to mount robust and durable responses with the potential to provide long-term protection from reinfection, and from emerging viral variants, remains to be understood. Insights into the strengths and limitations of the immune response in those who have had a successful outcome of natural infection can inform the future optimisation of vaccines. It is also crucial to understand the nature of the immune protection afforded to previously infected individuals whilst they await vaccination, especially with the ongoing delays in rollout and the lag in provision to low and middle-income countries.

Antibodies, in particular the neutralising fraction, provide a vital frontline defence to achieve protective immunity against viruses. An initial waning of antibody titres is typically seen after resolution of an acute viral infection.^3,4^ In the case of some viruses, long-lived plasma cells are then able to maintain antibodies for decades.^5–7^ By contrast, in the months following infection with other viruses, including human coronaviruses like SARS-CoV-2, neutralising antibodies continue to wane and can drop below the threshold of detection in a proportion of individuals.^3,8–13^ Even if antibodies are maintained, they may fail to provide sufficient functional flexibility to cross-recognise viral variants.^14–16^ However, antibody responses of inadequate titre or unable to cross-recognise variants can be compensated by a second line of defence provided by antigen-specific memory B cells (MBC), that are poised to react rapidly upon pathogen re-encounter.^17–19^ Not only can MBC provide a faster response on re-exposure to the virus, they are also able to diversify in the face of a mutating virus, resulting in more potent, affinity-matured antibody response and enhanced resistance to viral mutations.^9,20^

In this study, we therefore analysed whether MBC develop in elderly subjects following the resolution of SARS-CoV-2 infection and whether they can maintain functionality once neutralising antibodies (nAb) have waned. To address these questions, we studied elderly residents that had recovered from SARS-CoV-2 infection following outbreaks in three care homes in the UK, who had mild or asymptomatic infection, a substantial proportion of whom lost detectable nAb by five months after outbreak resolution. MBC were compared between the elderly care home residents and younger staff to assess the impact of ageing. We identified MBC specific for SARS-CoV-2 spike and RBD that persisted when serum nAb had completely waned. Their frequency, phenotype, isotype and function were analysed according to age and/or nAb loss, to inform the assessment and boosting of durable immunity in the elderly.

## Results

### SARS-CoV2 spike and RBD-specific memory B cells can persist after loss of neutralising antibodies

To study the role of MBC, we obtained PBMC from a subset (n=32) of a large cohort who survived COVID-19 with mild/asymptomatic infection after outbreaks in three care homes in April 2020 (Methods and Table S1).^21,22^ The care home cohort subset was selected to have a wide range of nAb titres detectable against live virus at the first sampling timepoint (T1, May 2020, Fig.1). By end September 2020 (T2, five months), 32% of all participants sampled had stable or increasing nAb to live virus. In contrast, 22% had declining titres, and 38% had lost detectable nAb (Fig.1a,b).

**Figure 1.**
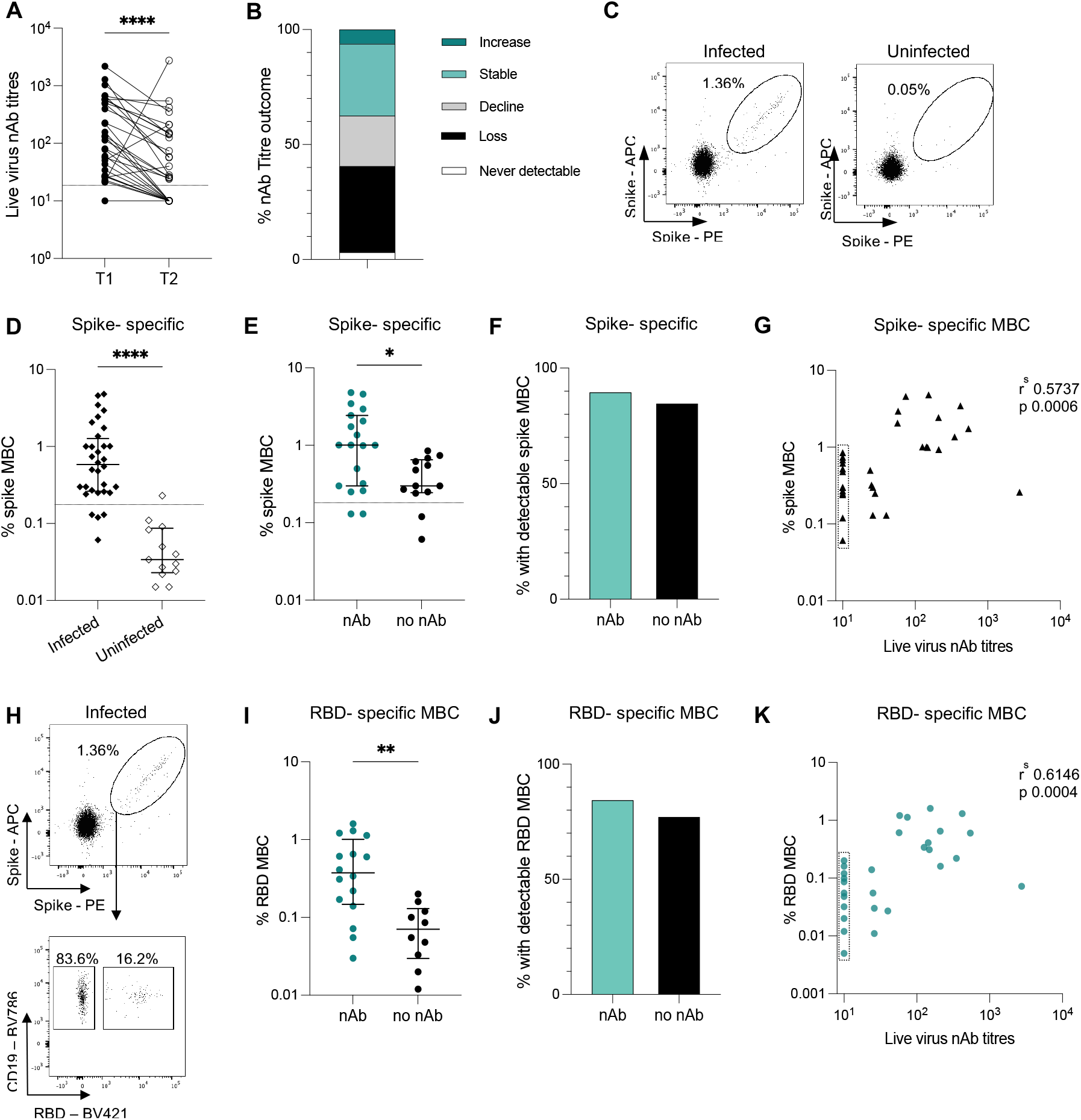
Spike and RBD-specific memory B cells persist five months post-SARS-CoV-2 infection despite waning neutralising antibodies. (**A**) Paired live virus nAb titres (nAb) at T1 and T2 of individuals infected prior to T1 (n=32). (**B**) Proportion of infected individuals with change in nAb indicated between T1 and T2: increase = ≥4 fold rise in nAb; static = >4 fold decrease <4 fold increase; decline ≥4 fold decrease; loss = no detectable nAb titres at T2 from detectable nAb at T1; never detectable = absence of nAb titres at T1 and T2 (n=32). (**C**) Representative FACS plots of dual staining with SARS-CoV-2 spike tetramers on MBC (CD3-CD14-CD19+CD20+CD38(+/-) IgD-excluding naïve (CD21+CD27-)) for previously infected (left) and uninfected (right) individuals. (**D, E**) Frequency of dual spike-specific MBC (**D**) in infected (n=32) and uninfected (n=13) and (**E**) in infected individuals with (nAb, n=19) and without (no nAb, n=13) detectable nAb at T2. Dashed lines indicate threshold for spike-specific responses determined by uninfected controls (Supplementary figure 1b). (**F**) Proportion of infected individuals with detectable spike-MBC above the threshold stratified by presence (nAb, n=19) and absence (no nAb, n=13) of detectable nAb at T2. (**G**) Correlation between frequency of spike MBC and live virus nAb titres in infected individuals (n=32). (**H**) Representative FACS plots of dual staining with SARS-CoV-2 spike tetramers on MBC (top panel) and RBD tetramer on dual spike specific cells (lower panel) of an infected individual. Minimum number of cells in spike-specific gate required for RBD probe analysis = 20 (**I**) Frequency of RBD-specific MBC in infected individuals with detectable spike-specific responses stratified by presence (nAb, n=16) and absence (no nAb, n=10) detectable nAb at T2. (**J**) Proportion of infected individuals with detectable RBD MBC stratified by presence (nAb, n=16) and absence (no nAb, n=10) of detectable nAb at T2. (**K**) Correlation between frequency of RBD MBC and live virus nAb titres in all infected individuals (n=29). (**A**) Wilcoxon matched pairs, p ≤0.0001. (**D, E, I**) Bars indicate median and interquartile range; Mann Whitney U Test; (**D**) p ≤0.0001, (**E**) p=0.0114, (**I**) p=0.0012. (**F, J**) Fisher’s exact test; (**F**) p= 0.6285, (**J**) p= 0.6664. (**G, K**) Dotted box indicates individuals with discordant MBC and nAb response. Spearman’s rank correlation. nAb, neutralising antibody; MBC, memory B cell, RBD, receptor binding domain. Analysis of RBD specific MBC only in those with ≥20 cells in spike-specific gate.

To compare MBC frequencies in those who had maintained or lost nAb, we stained PBMC with SARS-CoV-2 spike trimer tetramers, made by pre-incubating recombinant biotinylated trimeric spike protein with fluorescently-conjugated streptavadin.^15^ Dual staining with spike tetramers with two distinct fluorochromes was used to enhance the discrimination of true antigen-specific MBC (Fig.1c), as described previously.^23–25^ Frequencies of antigen-specific responses were calculated within the memory fraction of B cells (CD19^+^CD20^+^ excluding IgD^+^, CD38^hi^ and CD21^+^CD27^-^ naïve fractions, gating strategy in Fig.S1a, as previously described^26^). A threshold for background non-specific staining was set at mean+ 2SD of staining seen in an uninfected control cohort derived from the same care homes (seronegative at both time points, Table S1) and from pre-pandemic healthy donor samples (Fig.S1b).

Spike-specific MBC were detectable in 28 of the 32 tested 5 months post-infection (Fig.1d). The frequency of spike-specific MBC was reduced in those who had lost nAb compared to those in whom they were still detectable (Fig.1e). Of note, however, most of those (85%) who had lost detectable nAb still had some persistent spike-specific MBC, a comparable proportion to that in the group maintaining nAb (Fig.1f). The frequency of spike-specific MBC correlated significantly with the strength of the nAb response (nAb titre to live virus) at 5 months (Fig.1g); however, there was partial discordance due to detection of spike-specific MBC in most individuals with no nAb (dotted box, Fig.1g).

Next, we analysed the MBC response specifically directed against RBD since this is the region within spike which many SARS-CoV-2-specific nAb target.^15,27–29^ RBD-specific MBC were identified by gating on dual spike tetramer-staining populations that also stained with a tetramer formed from recombinant biotinylated RBD protein pre-incubated with fluorescently-conjugated streptavidin (Fig.1h). RBD-specific responses were detectable in 26 of the 28 with sufficient magnitude spike-specific MBC responses (>20 dual-spike+ cells) to allow analysis of the RBD-co-staining cells (Fig.1i). The frequency of RBD-specific MBC was significantly reduced in the group who had lost nAb compared to those with stable (or waning but still detectable) nAb (Fig.1i). However, as noted with spike-specific MBC, some RBD-specific MBC remained detectable in most of the cohort, irrespective of whether or not they had lost nAb (Fig.1j). Overall, the magnitude of RBD-specific MBC correlated with nAb titres, although again there was some discordance due to RBD MBC in those who had lost nAb (dotted box in Fig.1k). Importantly, both the RBD positive and RBD negative components of the spike-specific B cell response significantly correlated with nAb titres, though slightly more robustly for the RBD positive subset (Fig.1k, Fig.S1c). This highlights the importance of the RBD as the major target for neutralising antibodies, whilst also underscoring the contribution of antibodies targeting regions outside of the RBD (for example the NTD of the spike protein^15,29–31^) to the neutralising antibody response, at the 5 month timepoint in this cohort.

These data therefore revealed the persistence of detectable, albeit reduced, MBC specific for both spike and RBD in most people whose nAb titres against live virus had fallen below the threshold of detection. Thus, loss of detectable nAb 5 months after asymptomatic/mild infection is frequently compensated by the presence of a memory response primed to respond upon re-exposure.

### Comparable persistence of spike and RBD-specific MBC in elderly care home residents and younger staff

The care home cohort was constructed to sample two comparator groups: elderly residents (median age 86yrs, range 66-96) and a control group of younger staff (median age 56yrs, range 41-65). Five months after asymptomatic/mild infection, similar proportions of staff and residents had lost detectable nAb (Fig.2a), and those who maintained them had similar titres (Fig.2b). We postulated that there may, nevertheless, be a defect in the maintenance of spike/RBD-specific MBC in the elderly compared to younger age group. However, spike-specific MBC were maintained at similar frequencies and in comparable proportions of the elderly residents and younger staff (Fig.2c,d). There were no clear trends for spike-specific MBC to decrease with increasing age, even in residents in their nineties (Fig.2e).

**Figure 2.**
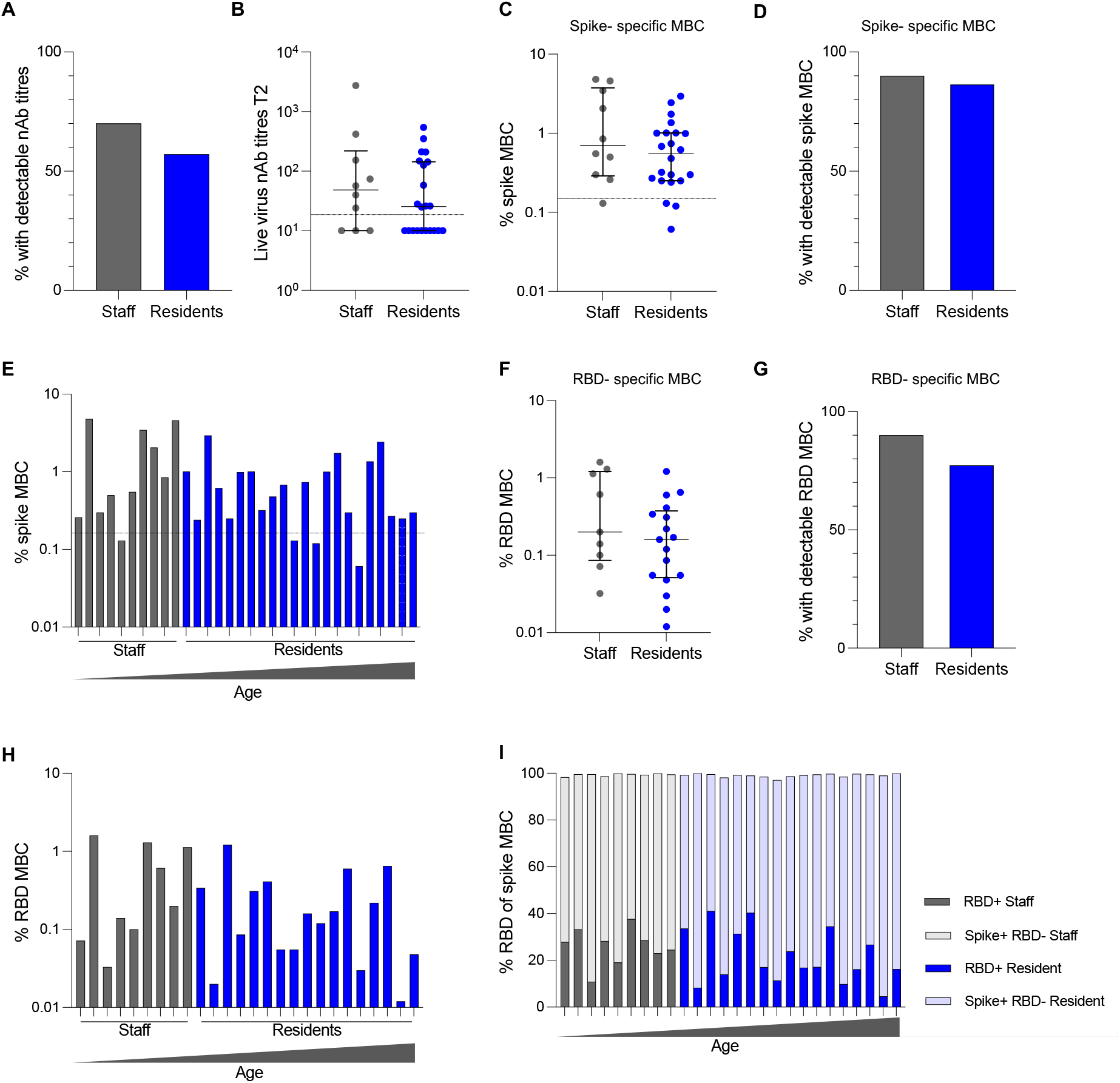
Comparable persistence of spike and RBD-specific memory B cells in elderly care home residents and younger staff. (**A**) Proportion of staff (n= 10) and residents (n=21) with detectable nAb at T1 who continued to have detectable nAb at T2. (**B**) nAb titres at T2 for all infected individuals stratified by staff (n=10) and residents (n=22). Dashed line indicates assay threshold for detection, undetectable titres assigned a value of 10. (**C**) Frequency of dual spike-specific MBC in staff (n=10) and residents (n=22). (**D**) Proportion of infected individuals with detectable spike-MBC stratified by staff (n=10) and resident (n=22) status. (**E**) Frequency of dual spike-specific MBC for staff (grey) and residents (blue) ordered by age from youngest on the left to oldest on the right. (**F**) Frequency of RBD-specific MBC in staff (n=9) and residents (n=17) with detectable spike specific responses. (**G**) Proportion of infected individuals with detectable RBD MBC stratified by staff (n=10) and resident (n=22) status. (**H)** Frequency of RBD-specific MBC for staff (grey) and residents (blue) ordered by age from youngest on the left to oldest on the right. (**I**) Proportion of dual spike specific cells with specificity for RBD (staff= dark grey; residents = dark blue), or non-RBD region (staff = pale grey; residents = pale blue) in staff (n=9) and residents (n=17). (**A, D, G**) Fisher’s exact test; (**A**) p>0.9999, (**D**) p> 0.9999, (**G**) p= 0.6367. (**B, C, F**) Bars indicate median and interquartile range; Mann Whitney U test; (**B**) p= 0.4367, (**C**) p= 0.2552, (**F**) p=0.2359. (**C, E**) Dashed line indicates threshold for spike-specific responses determined by uninfected controls (Supplementary figure 1b).

Similarly, RBD-specific MBC were equally well-maintained in the residents and staff (Fig.2f,g), with no decline in their frequencies (as a fraction of total MBC) with increasing age (Fig.2h). RBD-specific MBC comprised a variable proportion of the total spike-specific MBC response (4.6 to 41.0%; median 24.0%), the remainder representing B cells targeting non-RBD regions of spike. The proportions of RBD and non-RBD-binding spike-specific MBC again showed no changes with age (Fig.2i).

### Skewed isotype of spike/RBD-specific B cells associates with loss of neutralising antibodies

Having identified and quantified antigen-specific B cells with tetramer staining, we were able to apply high-dimensional multiparameter flow cytometry to phenotype these low frequency populations without any *in vitro* manipulation. We investigated the immunoglobulin isotype, memory phenotype, homing markers and transcription factor usage of spike and RBD-specific B cells, and global B cells (Fig.3).

**Figure 3.**
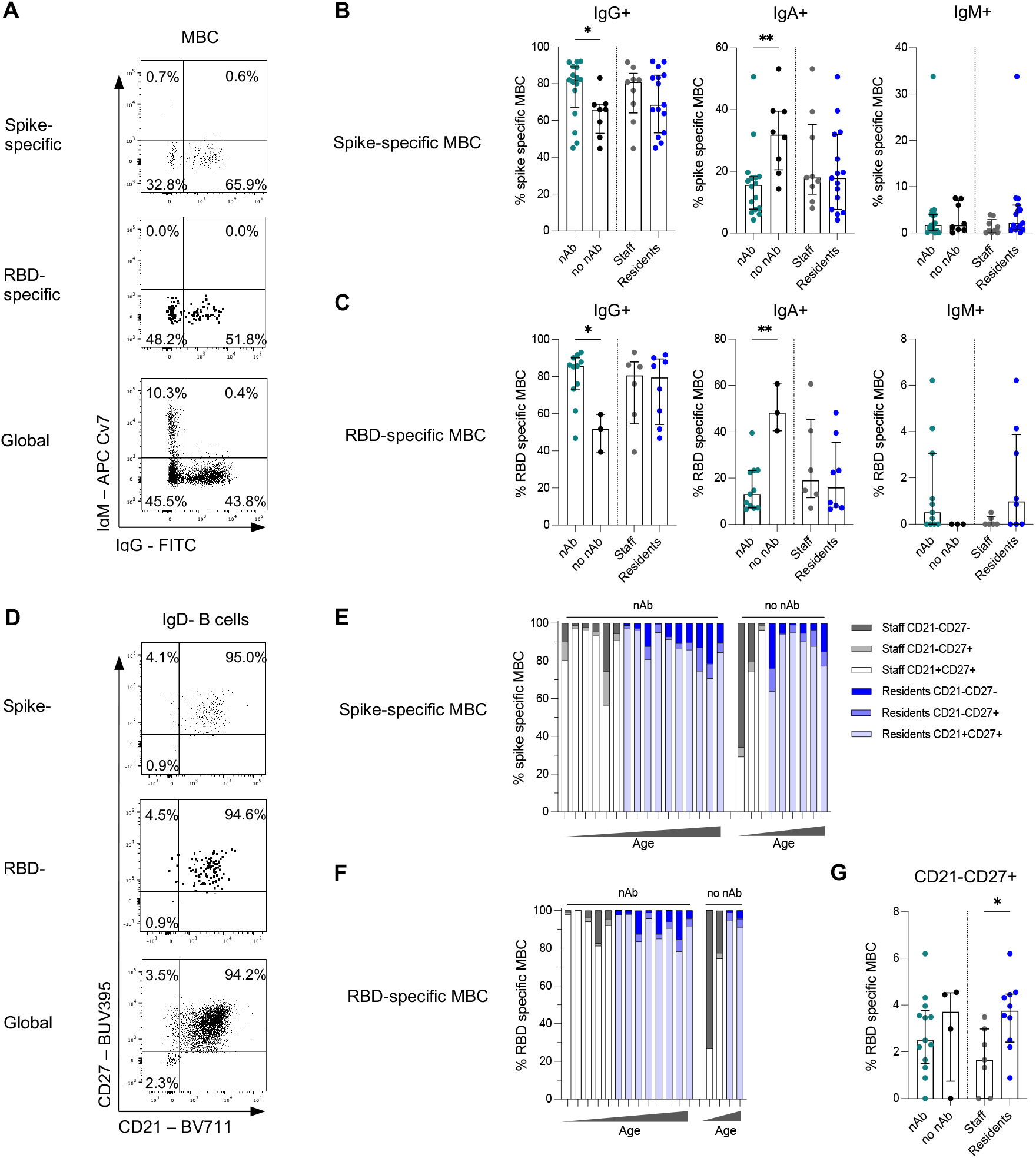
Preserved memory phenotype but skewed isotype of spike/RBD-specific B cells with loss of neutralising antibodies. (**A**) Representative FACS plots of IgM and IgG on spike-specific (top panel), RBD-specific (middle panel), and global (bottom panel) CD19+CD20+CD38lo/neg IgD-MBC from an infected individual. (**B**) Frequency of IgG+, IgA+ (denoted by IgD-, IgG-, IgM-) and IgM+ spike-specific MBC stratified by presence (nAb, n=16) and absence (no nAb, n=8) of detectable nAb at T2, and by staff (grey, n=9) and resident (blue, n=15) status. (**C**) Frequency of IgG+, IgA+ (denoted by IgD-, IgG-, IgM-) and IgM+ RBD-specific MBC stratified by stratified by presence (nAb, n=11) and absence (no nAb, n=3) of detectable nAb at T2, and by staff (grey, n=6) and resident (blue, n=8) status. (**D**) Representative FACS plots of CD21 and CD27 gating on spike-specific (top panel), RBD-specific (middle panel), and global (bottom panel) CD19+CD20+CD38lo/neg IgD-MBC from an infected individual. (**E**) Frequency of CD21-CD27+, CD21+CD27+ and CD21-CD27-MBC subsets of spike-specific MBC stratified by presence (nAb, n=16) and absence (no nAb, n=9) of detectable nAb at T2 ordered by increasing age. (**F**) Frequency of CD21-CD27+, CD21+CD27+ and CD21-CD27-MBC subsets of RBD-specific MBC stratified by presence (nAb, n=13) and absence (no nAb, n=4) of detectable nAb at T2 ordered by increasing age. (**G**) Frequency of CD21-CD27+ RBD-specific MBC stratified by presence (nAb, n=13) and absence (no nAb, n=4) of detectable nAb at T2, and by staff (grey, n=7) and resident (blue, n=10) status. (**B, C, G**) Bars indicate median and interquartile range; Mann Whitney U test; (**B**) IgG p=0.0382; ns, IgA p=0.0045; ns, IgM ns; ns; (**C**) IgG p=0.0220; ns, IgA p=0.0055; ns, IgM ns; ns; (**G**) ns; 0.0180. Analysis of individuals ≥50 cells in the relevant parent gate for all phenotypic analysis.

The vast majority of SARS-CoV-2 MBC expressed IgG, with a similar isotype distribution observed between spike and RBD-specific MBC (Fig.3a,b,c). However, individuals with persistent nAb had a higher frequency of IgG isotype expressing spike- and RBD-specific MBC than their counterparts who had lost nAb (Fig.3b,c), indicating the establishment of a robust, class-switched memory response in these individuals. In contrast, individuals whose nAb had waned below detectable limits had lost more IgG, and had a relative preservation of IgA class-switched spike- and RBD-specific MBC (Fig.3b,c). Elderly residents similarly showed a trend towards less IgG on their spike-specific MBC but, overall, no significant skewing of their immunoglobulin class-switching compared to younger staff (Fig.3b,c). Global B cells showed the same pattern of expression of different immunoglobulin isotypes on their surface in SARS-CoV-2 resolved donors as in uninfected controls, with roughly equal proportions of IgG and IgA and less than 15% IgM (Fig.S2a).

MBC subsets were examined using the combination of CD27 and CD21. The majority of spike and RBD-specific B cells had a classical resting memory phenotype (CD27^+^CD21^+^), characteristic of functional responses and comparable to the global MBC compartment, in both the elderly resident and staff groups (Fig.3d,e,f, Fig.S2b). ‘Double negative B cells’ have been associated with B cell dysfunction in ageing,^32–34^ and the ‘DN2’ subset with an extrafollicular short-lived plasmablast response in the acute phase of a cohort with severe COVID-19.^35^ However, at the five month timepoint following mild/asymptomatic infection in our cohort, neither the elderly nor those who had lost nAb showed any expansion of CD27^-^CD21^-^ B cells (Fig.3e,f) or the DN2 subset (CD27^-^CD21^-^CXCR5^lo^CD11c^hi^, Fig.S2c). Instead, there was a selective enrichment of the activated MBC subset (CD27^+^CD21^-^, previously described to be expanded in HIV and Ebola infection or after vaccination^36–38^) in the RBD-binding fraction in elderly residents, with the same trend in those who had lost nAb (Fig.3g). Those who had lost nAb also had reduced expression of the B cell homing molecules CXCR3 and CXCR5 on spike-specific and global MBC (non-significant trend and significant respectively, Fig.S2d,ef). T-bet, a transcription factor critical for acute antiviral function in B cells but associated with dysfunction in chronic infections and autoimmunity,^39–42^ also tended to be expressed at lower levels in the spike-specific MBC of those losing nAb (Fig.S2e,f).

Taken together, the isotype and memory phenotype of global and antigen-specific B cells was largely preserved in the elderly care home population, apart from an increase in spike-specific activated MBC. Individuals who maintained nAb had predominantly IgG-expressing antigen-specific MBC. In contrast, in those who had lost nAb by 5 months, whether staff or residents, residual antigen-specific B cells showed preferential preservation of IgA.

### Elderly residents maintain functional spike/RBD-specific B cells but at reduced frequency compared to younger care home staff

Having found that antigen-specific MBC could persist following complete waning of circulating nAb, we wanted to confirm their potential for functional recall upon re-encountering SARS-CoV-2. We therefore used cultured B cell ELISpots to examine the capacity of persistent SARS-CoV-2-specific MBC to differentiate into plasmablasts capable of secreting IgG capable of binding recombinant trimeric spike, S1 or RBD proteins.

ELISpots were performed using PBMC from 24 seropositive care home residents and staff, with the threshold for detection set at the highest observed value in an uninfected controls group (five seronegative care home residents and five pre-pandemic controls). Only individuals with responses detectable in a control total IgG well were included in analysis. Where responses were too numerous to count (TNTC), the highest number of spot-forming cells (SFCs) observed in the maximal response to the respective protein was used (Fig.S3a).

Functional recall responses to SARS-CoV-2 trimeric spike protein were observed in 21 of the 24 seropositive individuals tested, with ELISpots tending to be positive in more of those who had maintained nAb (Fig.4a). However, the majority of those who had lost detectable nAb still had a spike-specific response by ELISpot, with no significant difference in their magnitude compared to the nAb group (Fig.4a). ELISpots showed similar results for IgG binding S1 and RBD, with a trend to a lower proportion of positive results in those who had lost nAb but no significant difference in the magnitude of B cell recall responses in those maintaining serum nAb or not (Fig.4b,c).

**Figure 4.**
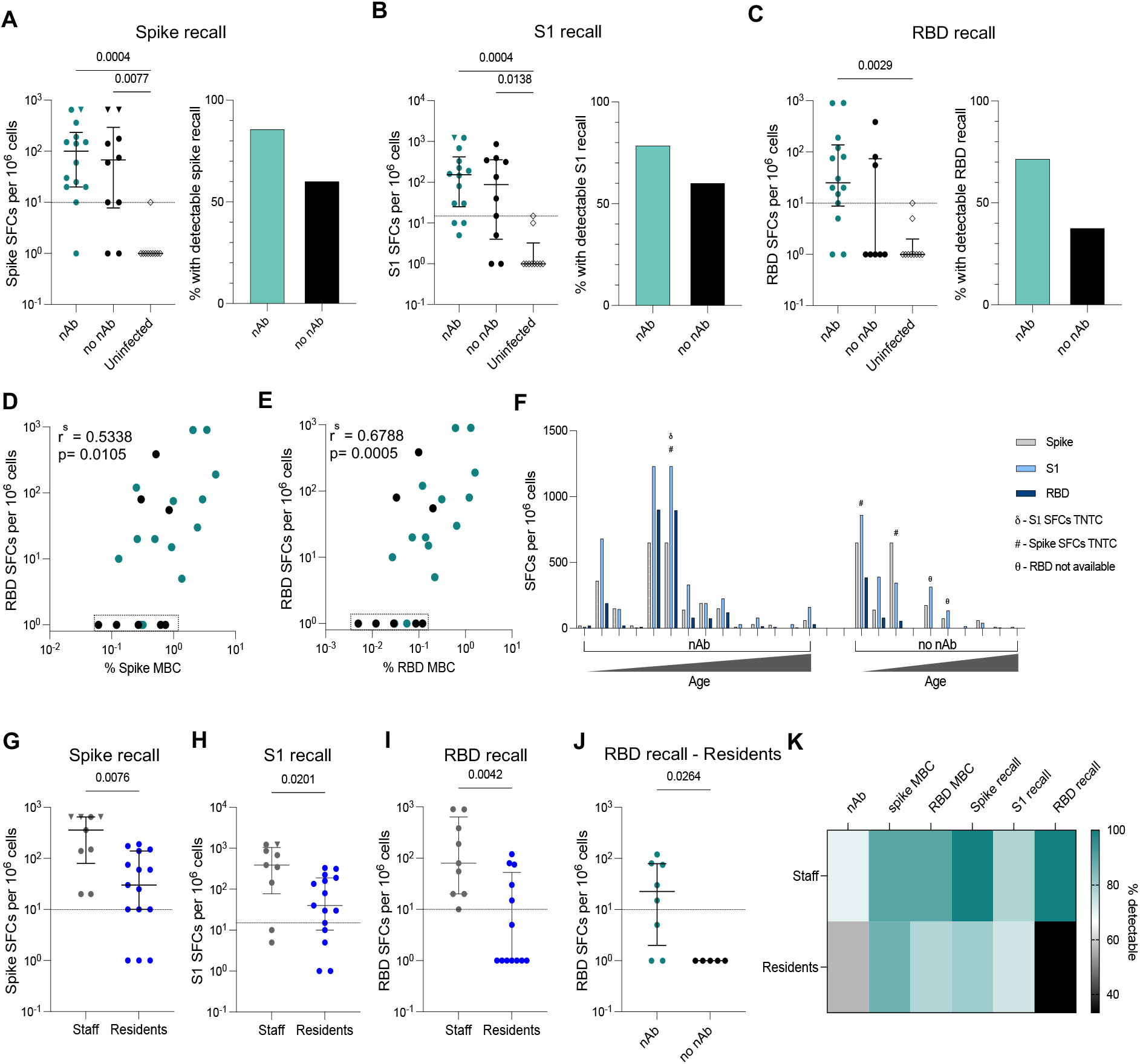
Elderly maintain some functional spike/RBD-specific B cells at reduced frequency compared to younger care home staff. (**A, B, C**) Left panels: SFCs per 1 million PBMC for infected individuals stratified by presence (nAb) and absence (no nAb) of detectable nAb at T2 and for uninfected controls. Right panel: proportion of infected individuals stratified by presence (nAb) and absence (no nAb) of detectable nAb at T2 with detectable recall responses. (**A**) Spike protein (nAb n=14, no nAb n=10, uninfected n=10), (**B**) S1 protein (nAb n=14, no nAb n=10, uninfected n=10), (**C**) RBD protein (nAb n=14, no nAb n=8, uninfected n=10). (**D, E**) Correlation between SFCs per 10^6^ PBMC to SARS-CoV-2 RBD protein and (**D**) frequency of spike specific MBC, (**E**) frequency of RBD positive MBC for those with nAb (green) and without nAb (black). (**F**) SFCs per 1 million PBMC to Spike (grey), S1 (pale blue), and RBD (dark blue) per individual stratified by presence (nAb, n=14) and absence (no nAb, n=10) of detectable nAb at T2 ordered by increasing age. (**G, H, I**) SFCs per 1 million PBMC for infected individuals stratified by staff and resident status, (**G**) Spike protein (staff n=9, resident n=15), (**H**) S1 protein (staff n=9, resident n=15, (**I**) RBD protein (staff n=9, resident n=13). (**J**) SFCs per 1 million PBMC to RBD protein for infected residents stratified by presence (nAb, n=8) and absence (no nAb, n=5) of detectable nAb at T2. (**K**) Summary heatmap of proportion of staff and residents with nAb titres detectable at T2, spike- and RBD-specific MBC by flow cytometry, and spike, S1 and RBD recall by ELISpot. **(A, B, C**) Left panels: Bars indicated median and interquartile range, dashed line indicates threshold indicated by seronegative and pre-pandemic controls. Kruskal Wallis multiple comparison ANOVA with Dunn’s correction, significance as indicated. **(A, B, C**) Right panels: Fisher’s exact test (**A**) p= 0.3413, (**B**) p= 0.3926, (**C**) p= 0.1870. (**D, E**) Dotted box indicates individuals with discordant MBC and ELISpot response. Spearman’s rank correlation. (**G, H, I, J**) Bars indicated median and interquartile range, dashed line indicates threshold indicated by seronegative and pre-pandemic controls. Mann Whitney U Test, significance as indicated. (**A, B, G, H**) Inverted triangle: individuals where the responses were TNTC, these individuals have been assigned the maximal response observed. (**F**) δ: SFCs TNTC in response to S1; #: SFCs were too numerous to count in response to Spike. For these individuals values have been assigned the maximum response observed on the plate for analyses. θ: RBD counts unavailable. Individuals with a zero response to any antigen have been assigned a value of 1 to allow plots to be drawn on a logarithmic scale. All statistical analysis performed using original values. SFC, spot forming cells; RBD, Receptor binding domain; MBC, memory B cells, TNTC, too numerous to count.

The magnitude of RBD recall response assessed by ELISpot showed a significant correlation with both spike and RBD MBC detection by tetramer staining (Fig.4d,e). However, there was some discordance due to individuals who had tetramer-binding spike or RBD B cells that did not produce detectable IgG by ELISpot (dotted boxes, Fig.4d,e), mainly in those who had lost nAb. Importantly, these data revealed that circulating antigen-specific B cells can be detected in the absence of functional recall.

Next, we compared functional responses to all three proteins for each individual, ranked according to nAb status and age. Individuals with strong recall to spike (as measured by ELISpot) tended to also have strong responses to S1 and RBD, whereas others had weak responses to all three antigens (Fig.4f). Functional MBC recall responses decreased with increasing age in both the groups, regardless of maintenance of serum nAb (Fig.4f). Thus, elderly residents had significantly lower ELISpot MBC responses against spike, S1, and particularly RBD, than the younger staff group (Fig.4g,h,i). Focusing on elderly residents who had lost nAb, we found that none of these individuals sustained MBC capable of functional recall to RBD (Fig.4j).

Overall, the measurement of nAb against live virus combined with assessment of spike and RBD-specific MBC by tetramer staining and functional ELISpot provided complementary insights into B cell immunity (Fig.4k). A substantial proportion of those who had lost neutralising activity against live virus, maintained spike and RBD-specific MBC detectable with one or both assays, regardless of age. However, some of those with persistent antigen-specific MBC could not mount a detectable functional response, particularly the elderly who had lost nAb (Fig.4k).

## Discussion

In this study we sampled a cohort of very elderly residents and younger staff who developed mild/asymptomatic SARS-CoV-2 infection during care home outbreaks, a high proportion of whom had lost nAb by five months. This allowed us to dissect the potential for B cell memory to persist beyond serum nAb, providing a back-up reserve to humoral immunity. We demonstrated that the majority of the cohort maintained detectable frequencies of spike and RBD-specific MBC by flow cytometry, even where they had lost circulating antibodies capable of live virus neutralisation. Tetramer staining allowed accurate *ex vivo* quantification and characterisation of antigen-specific MBC, revealing that individuals who had lost nAb had lower frequencies of spike and RBD-specific MBC, with a preserved classical memory phenotype but class-switching skewed away from IgG towards IgA. Elderly and younger recovered individuals infected in the same care home outbreaks maintained similar frequencies of spike and RBD-specific tetramer-staining B cells, with comparable phenotypes and isotypes. However functional assessment using ELISpot assays demonstrated that the persisting spike, and particularly RBD-specific, MBC had reduced potential for antibody production in the elderly.

The success of an infection or vaccine in inducing durable humoral immunity is dependent on the generation of long-lived plasma cells and MBC.^17–19^ The longevity of the plasma cell response, capable of sustaining antibodies, varies widely following different viral infections.^5–7^ A recent study has demonstrated the presence of bone marrow plasma cells secreting IgG against SARS-CoV-2 spike protein in fifteen of nineteen individuals examined seven months post infection,^43^ in line with the durability of some antibodies in the first year after mild infection. Nevertheless, many studies have also highlighted the potential for neutralising antibodies to SARS-CoV-2 to wane to a point where there is an, as yet ill-defined, risk of re-infection.^44–46^ Our study deliberately focuses on the role of MBC in those with waning or undetectable nAb to live virus, despite persistence of binding antibodies. MBC, previously identified in younger COVID-19 cohorts,^11,26,47,48^ can provide a crucial back-up by responding quickly to pathogen re-encounter or vaccination to form new plasmablasts, producing potent affinity-matured antibodies with more flexible recognition of viral variants;^9,20^ this is consistent with the enhanced nAb response described following vaccination of previously SARS-CoV-2 infected healthcare workers.^49^ Our demonstration that B cells of relevant specificities can still be detected even when nAb titres are waning or completely abrogated provides some reassurance that a memory response remains intact in the elderly. Future large-scale studies are needed to assess whether B cell memory serves as an independent correlate of protection or whether reliance on MBC to mount a new response in the absence of existing antibodies provides a critical window of opportunity for a virus that replicates as rapidly as SARS-CoV-2.

One strategy to combat antibodies that are waning or unable to cross-recognise emerging variants is the use of booster vaccines. Our finding that the elderly have impaired differentiation of their persistent spike/RBD-specific MBC into antibody producing cells detected by ELISpot assays provides biological rationale for a potential need for more frequent booster vaccination in this high-risk group. The frequency, phenotype and class-switching of antigen-specific B cells did not reveal obvious changes in the elderly group to account for this functional defect, other than an increase in the CD27^+^CD21^-^ subset. The activated CD27^+^CD21^-^ subset of MBC has recently been noted to remain expanded in some resolved COVID-19 patients,^50^ consistent with emerging literature supporting the possibility of prolonged antigen persistence, exemplified by a recent study detecting SARS-CoV-2 in the small bowel four months after asymptomatic infection.^9^ Our finding of more antigen-specific CD27^+^CD21^-^ MBC in the older age group raises the possibility there is more prolonged antigen persistence and resultant B cell activation following SARS-CoV-2 infection in the elderly. However, the ageing immune system is characterised by a tendency to low-level chronic inflammation,^51,52^ which could also contribute to prolonged activation of SARS-CoV-2 MBC. Analogous to our findings in elderly care home residents, both older subjects and those with HIV have been found to have persistent circulating MBC but defective plasmablast formation, resulting in reduced influenza vaccine-induced antibodies.^53,54^ Such age-related defects in B cell responses to vaccination have been attributed to a combination of B cell intrinsic senescence and defective T follicular helper cells (Tfh) in germinal centres.^55–57^

A caveat to our study is that we were only able to study circulating B cells, whilst additional recall responses may be compartmentalised within the mucosa. A recent study suggested mild infection can stimulate mucosal SARS-CoV-2-specific IgA secretion in the absence of circulating antibodies.^58^ The bias towards the retention of IgA+ spike/RBD-specific MBC in those who had lost all detectable serum nAb to live virus could therefore be reflective of a stronger mucosal response in these individuals. An increase in mucosal-homing IgA responses has been described as a feature of the ageing immune response,^59^ consistent with the older composition of our cohort. Alternatively, the relative preservation of IgA rather than IgG spike/RBD-specific MBC in those with the fastest waning nAb may simply reflect the recent observations that IgA dominates the early nAb response to SARS-CoV-2 infection,^56^ and may not decline as fast as the IgG response.^9,60^ Since our ELISpot assays did not measure the function of IgA isotype B cells, we may have under-estimated the full extent of residual SARS-CoV-2-specific responses, particularly in those with a more IgA-skewed response. In addition, several studies have shown that the magnitude of the MBC response to SARS-CoV-2 continues to increase beyond six months,^9,23,50,61^ again implying that we may have under-estimated the extent of recall potential in our cohort at five months. Future studies should also examine the preservation of non-spike-specific MBC with the potential to produce antibodies mediating antiviral effects beyond neutralisation, since other viral proteins (ORF3a, membrane and nucleocapsid) can play a dominant role in triggering antibody-dependent NK cell activation.^62^

In conclusion, by focusing on an elderly cohort with a high proportion of nAb loss, we demonstrated that this waning in the first line of humoral defence can be compensated by the presence of a reserve of adaptive B cell memory in the majority of cases. Our findings highlight the importance of including measures of B cell memory in larger studies of natural infection and vaccination to determine their role as additional correlates of protection. Our data underscore that identifying antigen-specific B cells by tetramer antigen staining is useful for quantitation and thorough *ex vivo* characterisation, but may not necessarily equate with the preservation of a functional response, as also observed in chronic viral infection.^42,63^ The relative preservation of IgA antigen-specific MBC in those with waned serum nAb raises the possibility that mucosal sequestered immunity may outlast that detectable in the circulation. Increased expansion of activated MBC in the elderly highlights the need to investigate whether they are more prone to prolonged stimulation from persistent reservoirs of SARS-CoV-2 antigen. A finding of concern was the lack of detectable functional recall to RBD in elderly donors who had lost nAb; given that RBD is the dominant site for nAb this supports the need for additional monitoring and/or booster vaccines to maintain sufficient antibodies to neutralise emerging variants in this highly vulnerable group.

## Materials and Methods

### Participants

SARS-CoV-2 antigen specific memory B cell (MBC) responses were compared between elderly care home resident and younger staff counterparts exposed to the virus within the same environment. Six care homes reporting SARS-CoV-2 outbreaks to Public Health England (PHE) were recruited to longitudinal SARS-CoV-2 RT-PCR and serological follow-up in April 2020 (T0).^1,21^ Serostatus of individuals within these homes at one month and five months after the outbreaks (T1 and T2 respectively) was established using binding and functional assays as previously described.^21,22^ Briefly, a native virus lysate assay (PHE) and/or receptor binding domain assay (RBD, PHE) determined seropositivity, and a live virus neutralising antibody assay to prototype England.2 SARS-CoV-2 virus was used to determine neutralising antibody titres.^21,22^

A total of 32 SARS-CoV-2 individuals (22 residents; 10 staff) were recruited, all of whom were seropositive by the binding assays described above at both sampling time points (T1: May 2020, T2: September 2020), alongside 11 SARS-CoV-2 seronegative control individuals from three of the care homes. Participants donated 30ml of blood to be processed for peripheral blood mononuclear cells (PBMCs) and serum five months after the initial outbreaks (T2). The investigation protocol was reviewed and approved by the PHE Research Ethics and Governance Group (REGG Ref NR0204). Written information regarding the study was provided to all participants; verbal consent for testing was obtained by care home managers from staff members and residents or their next of kin as appropriate.

Stored pre-pandemic samples from seven healthy individuals were used as controls, recruited under ethics number 11/LO/0421 approved by the ‘South East Coast - Brighton and Sussex Research Ethics Committee.

### Sample processing and data collection

Venepuncture blood samples collected in lithium heparin coated and serum separation tubes were used for isolation of PBMC and serum respectively. PBMC were isolated by density centrifugation using Pancoll human (PAN-Biotech). Isolated PBMC were frozen in foetal bovine serum (FBS) supplemented with 10% DMSO (Sigma Aldrich). Prior to use samples were thawed and washed in PBS. Serum was collected following centrifugation and stored at −80 degrees prior to use.

Clinical and laboratory data including age, gender, symptom status at T0 and SARS-CoV-2 RT-PCR status at T0 were available for all participants.(Table S1)^1^

### Protein expression and purification

Recombinant spike (S) and spike receptor binding domain (RBD) proteins of SARS-CoV-2 for antigen-specific B cell flow cytometry and ELISpot were expressed and purified as previously described.^15^ Briefly, spike glycoprotein trimer (uncleaved spike stabilised in the prefusion conformation (GGGG substitution at furin cleavage site and 2P mutation)^64^ and RBD protein^12^ were cloned into a pHLsec vector containing Avi and 6xHis tags. Biotinylated Spike and RBD were expressed in Expi293F cells (Thermofisher Scientific). Supernatants were harvested after 7 days and purified. For the production of biotinylated protein, spike and RBD encoding plasmids were co-transfected with BirA and PEI-Max in the presence of 200uM biotin.

Recombinant S1 protein constructs spanning SARS-CoV-2 residues 1-530 for ELISpot were produced as previously described.^28,30^ Briefly, codon-optimised DNA fragments were cloned into mammalian expression vector pQ-3C-2xStrep to create plasmids, which were then transfected into Expi293F cells growing at 37 in 5% CO2 atmosphere using ExpiFectamine reagent (Thermofisher Scientific). Proteins were purified by strep-tag affinity and subsequently size exclusion chromatography.

### Flow cytometry

High dimensional multiparameter flow cytometry was used for *ex vivo* identification of spike and RBD- specific B cells. Two panels (surface and intranuclear) of monoclonal antibodies (mAbs) were used to phenotype global and antigen specific subsets (Table S2). Biotinylated tetrameric spike (1ug) and RBD (0.5ug) were fluorochrome linked for flow cytometry by incubating with streptavidin conjugated APC (Prozyme) and PE (Prozyme) (Spike), and BV421 (Biolegend) (RBD) for 30 minutes in the dark on ice.

PBMC were thawed and incubated with Live/Dead fixable dead cell stain (UV, ThermoFisher Scientific) and saturating concentrations of phenotyping mAbs (Table S2) diluted in 50% 1xPBS 50% Brilliant Violet Buffer (BD Biosciences). For identification of SARS-CoV-2 antigen specific B cells 1ug per 500ul of stain each of tetrameric Spike-APC and Spike-PE and 0.5ug per 500ul stain of tetrameric RBD-BV421 were added to the cell preparation. Parallel samples stained with an identical panel of mAbs, but excluding the SARS-CoV-2 proteins (fluorescence minus one controls (FMO)) were used as controls for non-specific binding.

Cells were incubated in the staining solution for 30 minutes at room temperature, washed with PBS, and subsequently fixed with either fixation and permeabilization solution (BD Biosciences) or FoxP3 Buffer Set (BD Biosciences) according to the manufacturer’s instructions for surface and intranuclear staining respectively. Saturating concentrations of mAbs diluted in 1xPBS were added following permeabilization for the detection of intranuclear proteins. All samples were acquired on a Fortessa-X20 (BD Biosciences) and analysed using FlowJo (TreeStar).

B cell subsets were defined as follows: MBC - CD19^+^CD20^+^ excluding IgD^+^, CD38^hi^ and CD21^+^CD27^-^ naïve fractions, (gating strategy in Supp.Fig1a), DN2 - CD19^+^CD20^+^CD38^+/-^CD21^-^ CD27^-^CD11c^hi^CXCR5^lo^. For analysis of RBD-co-staining cells sufficient magnitude spike-specific MBC (≥20 dual-spike+ cells) were required. For phenotypic analysis of spike-specific and RBD-specific cells sufficient magnitude responses (≥50 cells in the relevant parent gate) were required.

### MBC recall response to SARS-CoV-2: ELISpot

To activate MBC differentiation, 1×10^6^ PBMC were stimulated with 1ug/ml R848 (TLR7/8 agonist; Resiquimod (Invivogen)) diluted in complete RPMI (cRPMI; RPMI supplemented with 10% FBS plus recombinant human IL-2 (20IU/ml; Peprotech), as previously described.^65,66^ Activated cells were incubated for six days with a media change on day three.

ELISpot plates (Mabtech) were pre-coated with recombinant SARS-CoV-2 trimeric spike (1μg/ml), S1 (1μg/ml) and RBD (10μg/ml) and anti-human IgG (1μg/ml Jackson Immunoresearch) overnight at 4°C. Coated plates were blocked with cRPMI with 10% FBS prior to the addition of cells. Cultured PBMC were added at varying concentrations depending on SARS-CoV-2 antigen and incubated at 37°C at 5% CO2 for 18 hours: 50,000 cells/ well to detect spike-specific IgG secreting cells; 100,000 cells/well to detect S1 and RBD IgG secreting cells; and 1000 cells/well for detection of total IgG secreting cells. To control for non-specific binding, uncoated control wells were incubated with 100,000 pre-stimulated cells. The following day, ELISpot plates were washed in filtered PBS supplemented with 0.5% Tween 20 (Merck) and incubated for four hours in the dark at room temperature with 1ug/ml goat anti-human IgG horse radish peroxidase antibody (Jackson Immunoresearch). Cells were again washed three times with PBS-Tween 20 (0.5%) and three times with PBS, then developed with 3-Amino-9-ethylcarbazole (AEC) substrate (BD Biosciences) according to the manufacturer’s instructions. ELISpot plates were washed with ddH20 before analysis using ViruSpot (Autoimmun Diagnostika). All conditions were performed in duplicate and responses averaged.

### Data analysis and statistics

Data were analysed using GraphPad Prism. Descriptive statistical analyses were performed. Continuous data that did not follow a normal distribution were described as medians with interquartile ranges and differences compared using the Mann-Whitney U test, Wilcoxon’s paired *t*-test or Kruskal Wallis test with Dunn’s post hoc test for pairwise multiple comparisons as appropriate. Contingency table analyses were conducted using Fisher’s exact test. Correlations for non-parametric data were assessed using Spearman’s rank correlation with 95% CI.

## Supplementary Materials

Fig. S1. **Gating strategy and threshold for detection of spike-specific responses**

Fig. S2. **Phenotyping of spike-specific and global MBC**

Fig. S3. **Representative ELISpot responses to SARS-CoV-2 proteins**

Table S1. **Cohort Characteristics**

Table S2. **Antibody list**

## Acknowledgements

The authors are very grateful to the care home managers, staff and residents; without their support and engagement this investigation would not have been possible. The authors would also like to thank the staff in the Immunisation and Countermeasures Department, in particular Maria Zavala, the Virus Reference Department, PHE Operations, the London Coronavirus Response Cell, in particular Nalini Iyanger and Jonathan Fok, PHE Field services, and the Maini laboratory for their help coordinating this investigation. We thank Peter Cherepanov of the Francis Crick Institute for supplying recombinant S1 antigen.

## Author contributions

Conceptualization: AJS, MZ, LEM, MKM; Methodology: AJS, ARB, LM, KJD, SNL, LEM, MKM; Investigation: AJS, ARB, SL, MP, RG, CRS, LEM; Sample and clinical data acquisition: AJS, FA, SNL, JYC; Analysis: AJS, ARB, SL, MP, RG, LEM, MKM; Funding acquisition: SNL, JYC, MZ, MKM; Supervision: SNL, MZ, LEM, MKM; Writing – original draft: AJS, MKM; Writing – review & editing: All authors

