## Supplementary Material for "SARS-CoV-2-specific memory B cells can persist in the elderly despite loss of neutralising antibodies"

#### Figure S1.

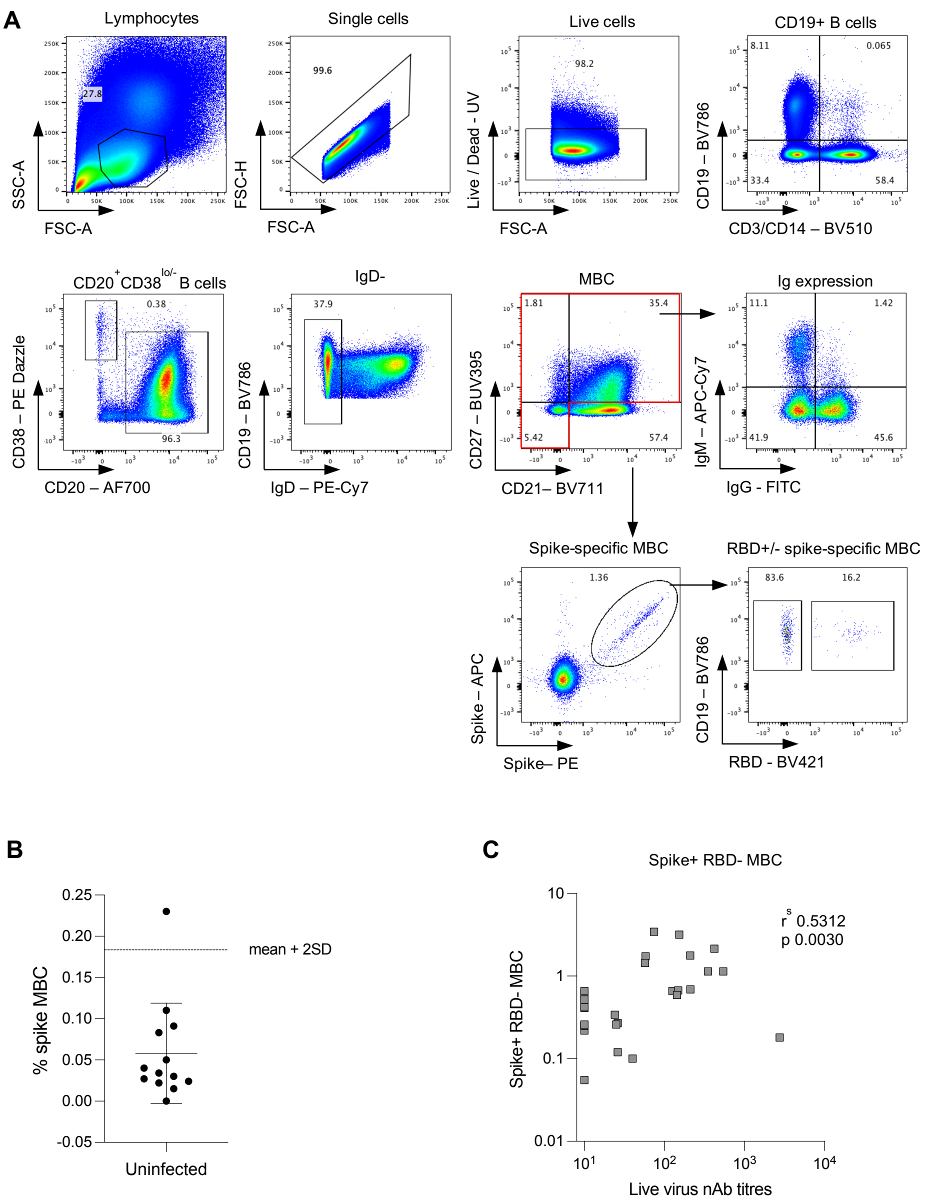


#### Gating strategy and threshold for detection of spike-specific responses

(**A**) Representative FACS plot showing gating strategy to determine populations indicated. Spike- and RBD-specific gating applied to IgD- MBC gate. (**B**) Frequency of spike-specific MBC in uninfected individuals used to determine threshold for detectable levels of spike-specific cells for uninfected individuals by calculating mean + 2SD. n=13 (n=11 uninfected, seronegative and n=2 pre-pandemic controls). (**C**) Correlation of frequency of spike-specific RBD- negative MBC and live virus nAb for all individuals with ≥20 cells in spike-specific gate (n=29). (**B**) Bars indicate mean and standard deviation. (**C**) Spearman's rank correlation. MBC – Memory B cell; Ig – immunoglobulin; RBD – receptor binding domain; nAb – neutralising antibody.

**Figure S2**.

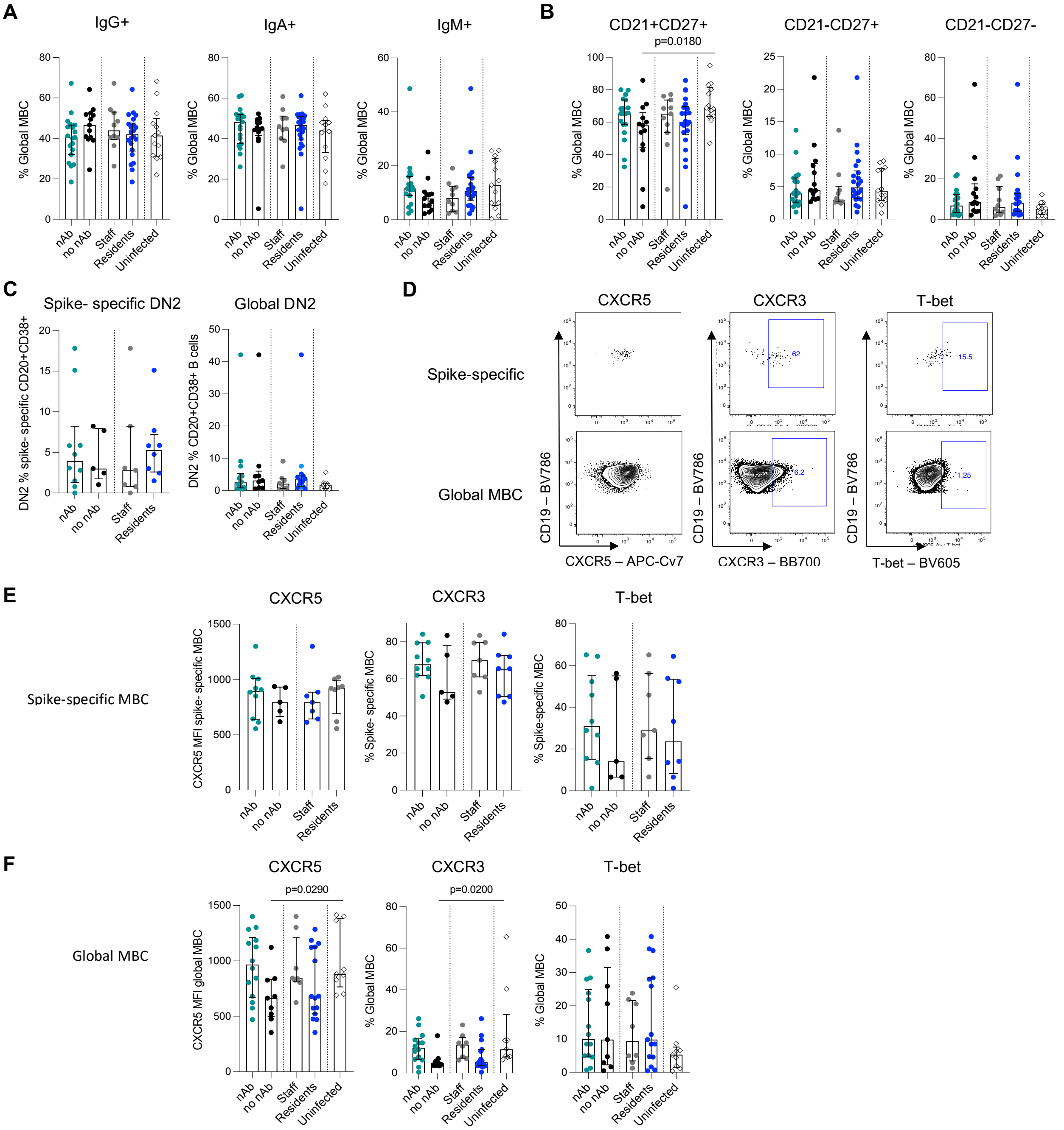


### Phenotyping of spike-specific and global MBC

**A**) Summary data of expression of immunoglobulin isotypes IgG, IgA and IgM on global MBC stratified by presence (nAb, n=19) and absence (no nAb, n=13) of detectable nAb at T2, and by staff (grey, n=10) and resident (blue, n=22) status, and uninfected controls (n=13). (**B**) Summary data of frequency CD21-CD27+, CD21+CD27+ and CD21-CD27- global MBC subsets stratified by presence (nAb, n=19) and absence (no nAb, n=13) of detectable nAb at T2, and by staff (grey, n=10) and resident (blue, n=22) status, and uninfected controls (n=13). (**C**) Left panel: Frequency of spike- specific cells with a DN2 phenotype ($CXCR5^{lo}$ $CD11c^{hi}$ CD21-CD27-) as a proportion of CD20+CD38+ B cells stratified by presence (nAb, n=10) and absence (no nAb, n=5) of detectable nAb at T2, and by staff (grey, n=7) and resident (blue, n=8) status. Right panel: Frequency of global CD20+CD38+ B cells with a DN2 phenotype stratified by presence (nAb, n=19) and absence (no nAb, n=13) of detectable nAb at T2, and by staff (grey, n=10) and resident (blue, n=22) status, and uninfected controls (n=13). (**D**) Representative FACS plots of CXCR5, CXCR3 and T-bet on spike- specific MBC (top panel) and global MBC (bottom panel). (**E**) Summary data of expression of CXCR5 (MFI), CXCR3 (frequency), and T-bet (frequency) of spike-specific MBC stratified by presence (nAb, n=10) and absence (no nAb, n=5) of detectable nAb at T2, and by staff (grey, n=7) and resident (blue, n=8) status. (**F**) Summary data of expression of CXCR5 (MFI), CXCR3 (frequency), and T-bet (frequency) of global MBC stratified by presence (nAb, n=19) and absence (no nAb, n=13) of detectable nAb at T2, and by staff (grey, n=10) and resident (blue, n=22) status, and uninfected controls (n=13). (**A, B, C, F**) Bars indicate median and interquartile range; Kruskal Wallis test with Dunn's post hoc analysis for multiple comparisons between nAb, no nAb and uninfected subgroups and staff, resident and uninfected subgroups respectively on global populations; significance where indicated. (**C, E**) Bars indicate median and interquartile range; Mann Whitney U test comparison between nAb and no nAb, and staff and resident subgroups respectively for spike-specific subsets; significance where indicated. Analysis of individuals ≥50 cells in the relevant parent gate for all phenotypic analysis.

**Figure S3.**

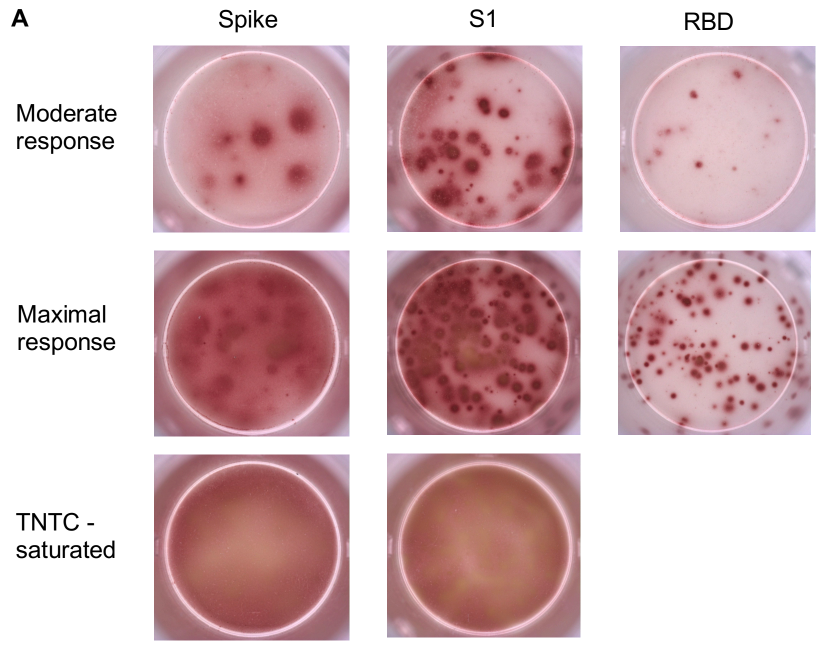


**Representative ELISpot responses to SARS-CoV-2 proteins**

(**A**) Representative ELISpot responses to SARS-CoV-2 Spike, S1, and RBD proteins. Top row demonstrates moderate response observed, middle row maximal responses, and bottom row saturated responses that were too numerous to count (TNTC). In the case of samples with saturated responses these individuals were assigned the value of the maximal response observed and highlighted as TNTC.

**Table S1.**

| | Residents | Staff | Uninfected |
|---|---|---|---|
| Number | 22 | 10 | 11 |
| Age years (median; IQR) | 86; 75-89 | 56; 50-60 | 82; 77-87 |
| Sex (n, (%) female) | 16 (72.7) | 7 (70.0) | 6 (54.5) |
| Symptom status T0* | 13 | 3 | 2 |
| PCR positive T0 | 13/21** | 5/9** | 0/11 |
| SARS-CoV-2 Seropositive T1*** | 22 | 10 | 0 |
| SARS-CoV-2 Seropositive T2*** | 22 | 10 | 0 |

**Cohort Characteristics**

* Symptom status assessed during the 14 days before and at T0 was collected for all staff, who self-reported any symptoms, and residents, whose symptoms were recorded the care home staff. Daily interviews were undertaken with individual care homes to identify any newly symptomatic individuals for the subsequent 14 days. Typical COVID-19 symptoms at that time included fever 37.8$^{o}$C, shortness of breath or cough, while atypical symptoms included, but were not restricted to, new confusion, reduced alertness, fatigue, lethargy, reduced mobility and diarrhoea.

** Two individuals (1 staff and 1 resident) did not have a nasal swab for SARS-CoV-2 RT-PCR taken at T0.

*** As assessed by native virus lysate assay (PHE) and / or receptor binding domain assay (RBD).

**Table S2.**

| Marker | Fluorochrome | Clone | Source | Cat. number |
|---|---|---|---|---|
| CD3 | BV510 | OKT3 | Biolegend | 317332 |
| CD11c | FITC | B-ly6 | BD | 561355 |
| CD13 | BV510 | M5E2 | Biolegend | 301842 |
| CD19 | BV786 | HIB19 | BD | 740968 |
| CD20 | AF700 | 2H7 | BD | 560631 |
| CD21 | BV711 | B-ly4 | BD | 563163 |
| CD24 | PE-CY7 | ML5 | Biolegend | 311120 |
| CD27 | BUV395 | L128 | BD | 563815 |
| CD38 | PE-CF594 | HIT2 | BD | 562288 |
| CD183 (CXCR3) | BB700 | 1C6/CXCR3 | BD | 566532 |
| CD185 (CXCR5) | APC-Cy7 | J252D4 | Biolegend | 356926 |
| IgD | PE-Cy7 | IA6-2 | BD | 561314 |
| IgG | FITC | G18-145 | BD | 560952 |
| IgM | APC-Cy7 | MHM-88 | Biolegend | 314520 |
| STREP | APC | - | Prozyme | PJ25S |
| STREP | PE | - | Prozyme | PJRS25 |
| STREP | BV421 | - | Biolegend | 405226 |
| T-bet | BV605 | 4B10 | Biolegend | 644817 |
| Viability | UV | - | Invitrogen / Thermofisher | L23105 |

**Antibody list**
